# Garden, greenhouse or climate chamber? Experimental conditions influence whether genetic differences are phenotypically expressed

**DOI:** 10.1101/2023.12.06.570376

**Authors:** Pascal Karitter, Martí March-Salas, Andreas Ensslin, Robert Rauschkolb, Sandrine Godefroid, Hendrik Poorter, Johannes F. Scheepens

## Abstract

1. Common-environment experiments are important to study genetically-based phenotypic variation within and among plant populations. Such experiments can be performed in an experimental garden, greenhouse or climate chamber. However, phenotypic expression may be strongly affected by the environmental conditions and influenced by parental and storage effects. Hence, it is unclear if results from common-environment experiments are reproducible across multiple experimental setups.
2. In this study, we assessed the effects of three different growth facilities – outdoor garden, greenhouse, and climate chamber –, on phenotypic expression. We compared ancestral and descendant genotypes of the same population of *Leontodon hispidus*. We also evaluated differences in phenotypic expression between plants grown after one (F1) vs. two (F2) intermediate generations.
3. We observed strong differences among plants growing in different growth facilities. Furthermore, we found that descendants had larger rosettes than ancestors only in the greenhouse and they flowered later than ancestors exclusively in the climate chamber. We did not find significant differences between intermediate generations within the growth facilities.
4. Overall, our study demonstrates that environmental variation among growth facilities can dictate the presence and magnitude of phenotypic differences. This implies that absence of evidence for phenotypic differences is not evidence of absence. Experimental systems should be carefully designed to provide meaningful conditions related to the research question. Finally, growing a second intermediate generation did not impact the genetic differences of ancestors and descendants within the facilities, supporting that only one intermediate generation may be sufficient to reduce detectable parental and storage effects.

## Introduction

Common-environment experiments have been a powerful approach for evolutionary biologists ever since the Swedish botanist Göte Turesson established this approach one century ago. The ability of a given genotype to express different phenotypes depending on the environment (i.e., phenotypic plasticity; Sultan, 2000) complicates studies on the genetic basis of phenotypic differences, since field observations cannot be used to unravel genetic from plastic effects on the phenotype. However, by growing plants of the same species from different habitats in the same developmental environment, Turesson (1922) was able to attribute phenotypic variation among plant populations to genetic differences and identify natural selection as the main driver. This concept was further developed to study local adaptation by transplanting individuals originating from different populations reciprocally among the habitats from which they were sampled (Clausen et al., 1940; Kawecki and Ebert, 2004). Common-environment experiments can even be used to study rapid evolution of plant populations to recent environmental changes by experimentally applying different selection pressures over multiple generations (Hill and Caballero, 1992) or by using the resurrection approach, in which seeds collected before a potential selection pressure are revived and compared to plants from recently collected of the same population (Franks et al., 2018). Comparisons of these phenotypes in a common environment can then uncover evolutionary changes that occurred between samplings (Franks et al., 2007).

It is important that the results of experimental approaches are reproducible and robust to ensure their validation and generalization, but this can be challenging (Drummond, 2009). Plant responses can be affected by the choice of the growth facility (i.e., outdoor garden, greenhouse or climate chamber; Poorter et al., 2016) or non-genetic biases such as parental effects (Latzel et al., 2023). For instance, Massonnet and colleagues (2010) studied leaf growth variables and other traits of three *Arabidopsis thaliana* genotypes in 10 different laboratories and found that modest variations in growing conditions such as temperature, light quality and the handling of the plants can induce significant differences in molecular profiles and phenotypes. Consequently, relatively small differences in the growth facility can lead to significant differences in plant responses and results can be strongly affected by the choice of the common environmental conditions. Poorter and colleagues (2012) described experimental outdoor gardens as being relatively close to natural conditions in the field. They characterized outdoor gardens as commonly having low spatial heterogeneity but high temporal variations in temperature, light and water supply, high chance of plant damage (hail, herbivory, late frost) and episodes of extreme conditions (e.g., high irradiance causing high temperatures, and low precipitation). According to Poorter and colleagues (2012), greenhouses normally provide more buffered conditions with heating systems to protect against frost, adjustable shading screens to counteract high irradiance, and with a more controlled water supply. However, air temperatures can peak depending on the ventilation system, and the greenhouse can substantially shade the plants due to structural elements (Cabrera-Bosquet et al., 2016). In climate chambers, researchers can control the experimental conditions as reliable as possible, which would be optimal for reproducibility, but the conditions can be quite artificial and deviate strongly from field conditions with more heterogeneous light and humidity distributions. Especially light can have strong vertical gradients and heterogeneous horizontal gradients (Poorter et al., 2012). Furthermore, climate chambers provide limited space which can greatly affect the possible sample size for larger species and thus, have a trade-off between control over the environment and statistical power. These differences among growth facilities may be important, as certain environmental conditions may not elicit phenotypic variation, even though genetic variation is present among genotypes. Accordingly, Stanton and colleagues (2000) found that, compared to near-optimal growing conditions, more stressful conditions, which better mimic natural conditions, tend to increase phenotypic variation among genotypes of wild mustard (*Sinapis arvensis*). Consequently, is unclear if the results from common-environment experiments are consistent throughout different experimental environments and can be accurately reproduced. If not, the consequence would be that we may miss evidence of relevant genetic variation or, in contrast, that we may detect genetic variation that has currently no adaptive relevance in the field.

Studies on genetic differentiation can also be confounded by non-genetic or random variability induced by parental effects or storage. Parental effects occur, when the parental phenotype affects the phenotype of their offspring irrespective of the genes that are inherited (Badyaev and Uller, 2009; Auge et al., 2017). Seed provisioning is one major parental effect that can have a big impact on the offspring, because the resource availability and general environmental conditions (e.g., light quantity, quality and duration) of the mother plant can determine the amount of resources for the seeds and consequently affect seedling establishment and early life history traits (Herman and Sultan, 2011). Other parental effects include hormone-driven effects on physiology of the seedling or epigenetic processes through passing on distinct DNA methylations or chromatin changes (Blödner et al., 2007; Jablonka and Raz, 2009; Elwell et al., 2011; Herman and Sultan, 2011; Richards et al., 2017). Even though parental effects can have ecological and evolutionary significance (Latzel et al., 2023), their influence can be a source of bias in studies on genetic differentiation when parental environmental conditions differ among genotypes. Furthermore, long storage periods can affect seed viability and even post-emergence traits (Franks et al., 2018). In experiments where plants from the same population but largely different generation are compared, so-called ‘resurrection experiments’, storage duration and conditions may well have been different. Consequently, phenotypes of seedlings may express varying plastic responses (Weis, 2018). These biases can be minimized by acclimating the experimental plants under common environmental conditions for one or more generations before the start of the experiment (Kawecki and Ebert, 2004). Parental effects caused by the environment usually disappear after one generation in a new environment (Agrawal, 2002; Gianoli, 2002), but have also been shown to persist over multiple generations (Wulff et al., 1999). Latzel (2015) recommends at least two intermediate generations before the start of an experiment, as it increases the chance of evening out epigenetic modifications. Still, the method of growing intermediate generations is not always implemented in common-environment studies as it is time– and labor-intensive, especially if the study plants do not flower in the first year (Bischoff and Müller-Schärer, 2010; Rauschkolb et al., 2023).

In this study, we investigated the consistency of phenotypic differences of a resurrection common-environment experiment among different growth facilities. Furthermore, we tested for differences between results from experimental plants grown after one vs. two intermediate (i.e., refresher) generations. Absence of any differences would suggest that one intermediate generation is sufficient to reduce detectable parental, epigenetic or storage effects. We worked with the perennial herb *Leontodon hispidus* and used seeds of ancestors sampled in 1995 and of descendants sampled in 2018 collected from the same population in a calcareous grassland in Belgium. We grew two intermediate generations (F1 and F2) from both ancestors and descendants under common conditions and cultivated them in three different growth facilities: in an outdoor garden, in a greenhouse and in a climate chamber. We measured eight traits regarding growth, leaf anatomy and flowering phenology to cover a wide spectrum of functional traits. We hypothesized that (1) genetically-based phenotypic differences across traits tend to occur inconsistently across substantially different growth facilities. (2) We expected that phenotypic differences in evolved traits between ancestors and descendants would be strongest in less-controlled conditions such as the outdoor garden and weakest in the more optimal and constant conditions in the climate chamber. Finally, we hypothesized that (3) one intermediate generation is not enough to sufficiently reduce non-genetic differences between ancestors and descendants, which may be attributed to the storage or parental effects of ancestors.

## Material and Methods

### Study species

*Leontodon hispidus* (Asteraceae) is a perennial herbaceous herb and typically flowers from June to October (Kühn and Klotz, 2002). It is self-incompatible and pollinated by insects (Kühn and Klotz, 2002). It is widespread throughout Europe and commonly found in calcareous grasslands. Seed material was collected from two temporal origins, 1995 (ancestors) and 2018 (descendants), from a single population in a dry calcareous grassland in a Belgian nature reserve (50°47’35″N, 5°40’25″E). The distance to the nearest neighboring population is approximately 2 km, likely preventing the majority of gene flow into the population. The staff of the Meise Botanic Garden (Belgium) collected the ancestral seeds for conservation purposes and efforts were made to represent the genetic diversity of the population by collecting from as many individuals as possible dispersed throughout the population. The seed material from an unknown number of mother plants was cleaned, bulked, dried at 15% relative humidity, and stored at –20 °C at the seed repository of the Meise Botanic Garden. In the summer of 2018, we revisited the population and collected seeds from 20 mother plants. These seeds were cleaned, bulked and then stored at 4 °C. Rauschkolb et al. (2022a) analyzed genomic relatedness and allelic richness among individuals within both temporal origins (ancestors and descendants) and found similar levels of relatedness without obvious kinship structure, which supports the comparability of the sampling procedures and confirms that sufficient seeds were collected.

### Experimental design

Both ancestral and descendant seeds were grown for two consecutive intermediate generations (F1 and F2). For the first intermediate generation (F1), we sowed 300 seeds from each temporal origin and selected 15 random individuals from each temporal origin which were randomly pollinated by hand in net cages to prevent unintentional cross-pollination (Rauschkolb et al., 2022b). We used the seeds from the F1 intermediate generation for the F2 generation and grew them under similar conditions, this time using bumblebees (Natupol Seeds, Koppert GmbH, Straelen, Germany) as pollinators. Ultimately, seven maternal lines from the F1 intermediate generation and 8 maternal lines from the F2 intermediate generation yielded sufficient seed material for both temporal origins. For each maternal line, we used 12 seedlings grown individually in black 1.5 L pots that were prepared as follows: In July 2022, we placed 12 pots for each maternal line with cultivation soil (Spezial Substrat Typ T1b, Hawita GmbH, Vechta, Germany) in the greenhouse and sowed three seeds into each pot. All pots were watered three times a week to maximum soil capacity.

After the seedlings had developed their first true leaf, we thinned them to a single individual per pot, with the seedling moved to the center of the pot. We measured the initial size as rosette diameter and divided the pots randomly into three groups with four individuals from each maternal line. Each group was grown in a different growth facility for the rest of the experiment: outdoor garden, greenhouse or climate chamber. In total, we used 360 plants for this experiment (3 growth facilities × 2 temporal origins × 2 generations × 7/8 maternal lines × 4 replicates). Pots in the garden were placed on gravel 2 m below a shading cloth to reduce radiation and temperature stress (Schattiergewebe 45%, Nitsch GmbH, Kreuztal, Germany). Plants were randomized every two weeks and watered weekly to soil capacity for the duration of the experiment. In the greenhouse, the plants were placed 1 m below lamps with a combination of two fluorescent tubes (Lumilux HO 80W-865, Berlin, Germany and Gro-Lux FH 80W, Sylvania, Erlangen, Germany). The lamps were programmed to switch on between 6 am and 20 pm (14 h photoperiod) whenever the natural light intensity went below 360 from µmol m^-2^ s^-1^ outside. To avoid extreme temperatures, sliding shutters and lamps were programmed activate once light intensity surpassed 1100 from µmol m^-2^ s^-1^ outside. The climate chamber (ThermoTec GmbH, Weilburg, Germany) was set to 14h-10h day-night cycle with 21°C during the day and 18°C during night to simulate the start of the growing season. Air humidity was set to a constant 60% and the plants were placed 1 meter below halogen lamps (Radium HRI-BT 400W/D Pro Daylight, Lampenwerk GmbH, Wipperfürth, Germany).

In each growth facility, we inserted four temperature and soil moisture loggers (TMS-4 logger, Tomst, Prague, Czechia) in the center of black 1.5 L pots with the same cultivation soil and positioned them randomly among the pots with plants. The loggers monitored soil temperature at 5 cm depth, soil surface temperature, air temperature at 5 cm above the soil as well as soil moisture every 15 min. We used this data to calculate mean values of all four loggers from each environment and to derive daily mean values for all parameters (Fig. 1, Appendix S1; see Supplemental Data with this article). Furthermore, we calculated mean values for soil temperature, soil surface temperature and air temperature over the course of the whole experimental period (Table 1). On average, temperatures were intermediate in the garden with 21.2 – 22.6 °C, highest in the greenhouse with 23.9 – 26.1 °C, and lowest in the climate chamber with 20.4 –21.2 °C (Table 1).

**Figure 1.**
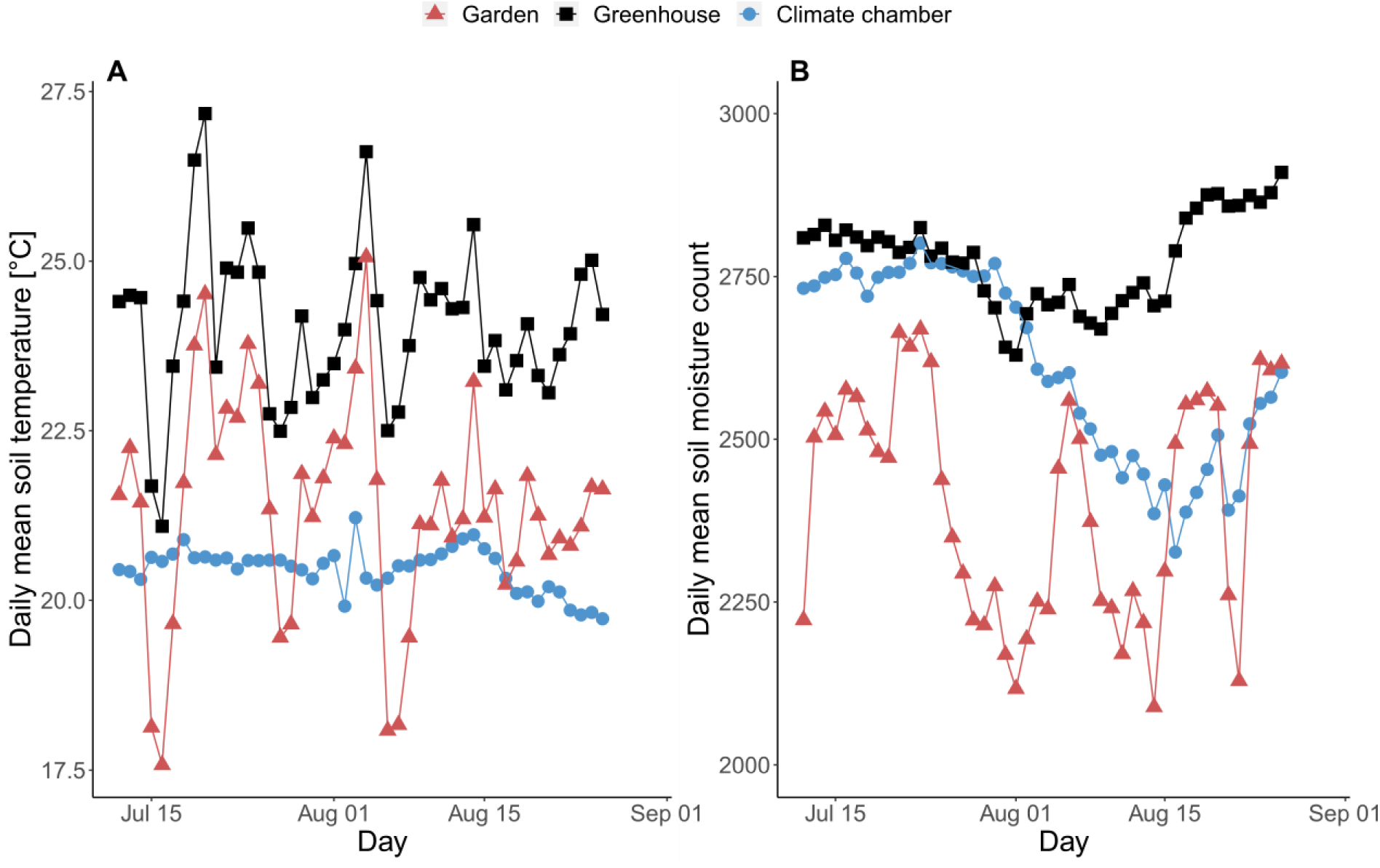
Daily mean soil temperature (A) and daily mean soil moisture count (B) of four random pots grown in different growth facilities. The growth facilities are garden (red triangles), greenhouse (black squares), and climate chamber (blue circles).

**Table 1.** Mean values and standard errors of environmental variables over the whole experimental period for all growth facilities.

| Growth facility | Temperature [°C] |  |  | Daily light integral<br>[mol·m<sup>-2</sup>·d<sup>-1</sup>] |
| --- | --- | --- | --- | --- |
|  | Soil (5cm) | Soil surface | Air |  |
| Garden | 21.2 ± 0.2 | 22.2 ± 0.3 | 22.6 ± 0.3 | 40.5 ± 1.6 |
| Greenhouse | 23.9 ± 0.2 | 25.5 ± 0.2 | 26.1 ± 0.2 | 7.5 ± 0.3 |
| Climate chamber | 20.4 ± 0.05 | 21.2 ± 0.03 | 20.7 ± 0.04 | 15.0 ± 1.0 |

At the start of the experiment at noon, we measured the light intensity using a light meter (Panlux electronic 2, GMC-Instruments, Nürnberg, Germany) in the outdoor garden and greenhouse at 12 spatially distributed pots at the same height 2 cm above the soil. After the end of the experiment, hourly solar radiation data (Wm^-2^) was extracted for the whole experimental period from a weather station in the outdoor garden (iMetos 1, Pessl Instruments GmbH, Weiz, Austria). To assess, how much radiation the plants received in the outdoor garden, we multiplied the solar radiation data by 0.55 to account for the shading cloth (45 % shading) and converted the data into Photosynthetic Photon Flux Density (PPFD, µmol•m^-2^•s^-1^) using the default function *Rg.to.PPFD()* from the *bigleaf* package (Knauer et al., 2018) in R (version 4.0.3, R Core Team, 2020). Fraction of incoming solar irradiance to photosynthetically active radiation (PAR) was set to 0.5 and the conversion factor was set to 4.6. According to the light intensity measurements at the start of the experiment, the greenhouse received 18.5 % of the light that the outdoor garden plants received. We used this ratio to approximate how much PPFD the plants in the greenhouse received over the course of the experiment by multiplying the PPFD data of the outdoor garden by 0.185. We summed all values per day to calculate the daily light integral (DLI) for each day and then calculated the mean DLI for the whole experimental period. For the climate chamber, we directly measured PPFD at 12 spatially distributed spots at pot height using a PAR sensor (PAR Special sensor SKP 210, Skye Instruments Ltd, Powys, UK). Since the photoperiod was constant in the climate chamber (14h), we multiplied the mean PPFD measurement (297.5 µmol•m^-2^•s^-1)^ by the photoperiod (in seconds) to calculate the DLI. The average DLI over the course of the experiment (Table 1) was highest in the garden (40.5 mol•m^-2^•d^-1^), lowest in the greenhouse (7.5 0 mol•m^-2^•d^-1^) and intermediate in the climate chamber (15.0 mol•m^-2^•d^-1^).

### Measurements

During the experiment, we recorded onset of flowering three times per week, because it is an essential trait in the life history of a plant and has often been observed to evolve under changing environmental conditions (Pelletier et al., 2009; Franks, 2015). We regarded a plant as flowering when the first anther was visible. Plants started flowering in August and after three months, most plants had flowered and we measured the rosette diameter as a measure of growth and ability to capture light. We also recorded the number of flowering stems as a proxy for reproductive success. For each individual, we measured the chlorophyll content of four randomly selected healthy and fully developed leaves in SPAD units using a chlorophyll meter (SPAD-502 Plus, Konica Minolta, Neu-Isenburg, Germany). We measured leaf area of three randomly selected healthy and fully developed leaves per plant with the smartphone application “easy leaf area free” (Easlon and Bloom, 2014). These leaves were dried in a dry oven at 60 °C for three days and then weighed together at a fine scale (CPA225D-0CE, e = 1 mg, Sartorius AG, Göttingen, Germany). At the same day, we harvested flower heads and flowering stems as reproductive biomass and leaves as vegetative biomass and dried these using the same procedure. This allows us to detect whether plants invest their resources into vegetative growth or rather into reproductive structures. In order to investigate responses in leaf anatomy, we calculated specific leaf area (SLA) by dividing the combined leaf area of the three selected leaves by their dry weight and calculated the leaf dry matter content (LDMC) by dividing the dry weight by the fresh weight. The weight of the three selected leaves was added to the vegetative biomass.

## Data analysis

All statistical analyses were performed using R (version 4.0.3, R Core Team, 2020). We used linear mixed effects models implemented in the *lme4* package (Bates et al., 2015) with temporal origin, generation, and environment as fixed factor, as well as their two– and three-way interactions. Maternal line was included as random factor and initial size as covariate. Rosette diameter, vegetative biomass, SLA, LDMC, onset of flowering, reproductive biomass, number of flowering stems and the SPAD measurements were used as response variables, and we applied appropriate transformations to these variables when necessary to improve normality and heteroscedasticity of model residuals (Table 2). All linear models were analysed using the function *Anova()*, and *P* values (Appendix S2) were adjusted for multiple testing with the method of False Discovery Rate (Benjamini and Hochberg, 1995) using the *fuzzySim* package (Barbosa, 2015). Tukey post-hoc tests were applied using the *emmeans* package (Lenth, 2021) whenever an explanatory factor with more than two levels was significant.

**Table 2.**
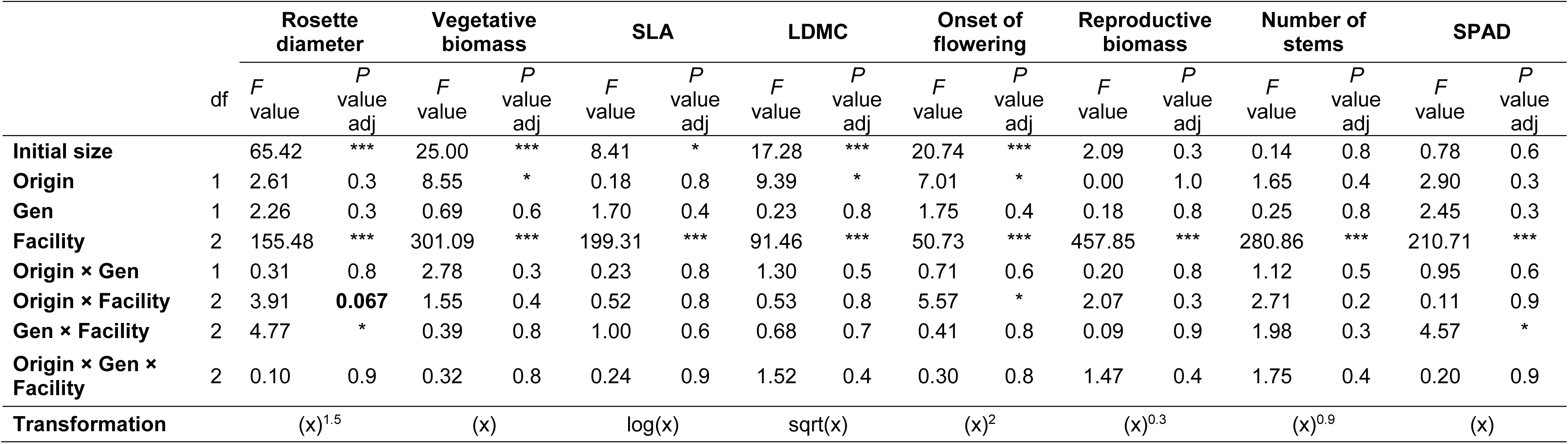
Results of the statistical models testing the effects of temporal origin (ancestors, descendants), intermediate generation (F1, F2), growth facility (garden, greenhouse, climate chamber) and their interactions on the response variables (y) rosette diameter, vegetative biomass, specific leaf area (SLA), leaf dry matter content (LDMC), onset of flowering, reproductive biomass, number of stems and SPAD measurements. We used linear mixed effects models with initial size as covariate and maternal line as random factor followed by ANOVA’s. Response variables were transformed if needed to fulfil parametric assumptions. Shown are degrees of freedom (*df*), *F* values and adjusted *p* values using False Discovery Rates. Significant *p* values are shown with asterisks (*: *P* ≤ 0.05; ***: *P* ≤ 0.001) and marginally significant values are shown in bold.

|  |  | Rosette diameter |  | Vegetative biomass |  | SLA |  | LDMC |  | Onset of flowering |  | Reproductive biomass |  | Number of stems |  | SPAD |  |
| --- | --- | --- | --- | --- | --- | --- | --- | --- | --- | --- | --- | --- | --- | --- | --- | --- | --- |
|  | df | *F* value | *P* value adj | *F* value | *P* value adj | *F* value | *P* value adj | *F* value | *P* value adj | *F* value | *P* value adj | *F* value | *P* value adj | *F* value | *P* value adj | *F* value | *P* value adj |
| Initial size |  | 65.42 | *** | 25.00 | *** | 8.41 | * | 17.28 | *** | 20.74 | *** | 2.09 | 0.3 | 0.14 | 0.8 | 0.78 | 0.6 |
| Origin | 1 | 2.61 | 0.3 | 8.55 | * | 0.18 | 0.8 | 9.39 | * | 7.01 | * | 0.00 | 1.0 | 1.65 | 0.4 | 2.90 | 0.3 |
| Gen | 1 | 2.26 | 0.3 | 0.69 | 0.6 | 1.70 | 0.4 | 0.23 | 0.8 | 1.75 | 0.4 | 0.18 | 0.8 | 0.25 | 0.8 | 2.45 | 0.3 |
| Facility | 2 | 155.48 | *** | 301.09 | *** | 199.31 | *** | 91.46 | *** | 50.73 | *** | 457.85 | *** | 280.86 | *** | 210.71 | *** |
| Origin × Gen | 1 | 0.31 | 0.8 | 2.78 | 0.3 | 0.23 | 0.8 | 1.30 | 0.5 | 0.71 | 0.6 | 0.20 | 0.8 | 1.12 | 0.5 | 0.95 | 0.6 |
| Origin × Facility | 2 | 3.91 | **0.067** | 1.55 | 0.4 | 0.52 | 0.8 | 0.53 | 0.8 | 5.57 | * | 2.07 | 0.3 | 2.71 | 0.2 | 0.11 | 0.9 |
| Gen × Facility | 2 | 4.77 | * | 0.39 | 0.8 | 1.00 | 0.6 | 0.68 | 0.7 | 0.41 | 0.8 | 0.09 | 0.9 | 1.98 | 0.3 | 4.57 | * |
| Origin × Gen × Facility | 2 | 0.10 | 0.9 | 0.32 | 0.8 | 0.24 | 0.9 | 1.52 | 0.4 | 0.30 | 0.8 | 1.47 | 0.4 | 1.75 | 0.4 | 0.20 | 0.9 |
| Transformation |  | (x)<sup>1.5</sup> |  | (x) |  | log(x) |  | sqrt(x) |  | (x)<sup>2</sup> |  | (x)<sup>0.3</sup> |  | (x)<sup>0.9</sup> |  | (x) |  |

## Results

The growth facility significantly affected all measured traits (Table 2). In the outside garden, plants had the lowest rosette diameter, and SLA, while those traits were highest in the greenhouse and intermediate in the climate chamber (Fig. 2AC). In contrast, vegetative biomass and LDMC were highest in the climate chamber followed by the garden, and lowest in the greenhouse (Fig. 2BD). A similar pattern can be observed in the onset of flowering (Fig. 3A): plants in the greenhouse and garden flowered at a similar time, but the onset of flowering of plants in the climate chamber was approximately 4 days delayed. The reproductive biomass, the number of flowering stems and SPAD values (Fig. 3BCD) were highest in the garden, intermediate in the climate chamber and markedly low in the greenhouse.

**Figure 2.**
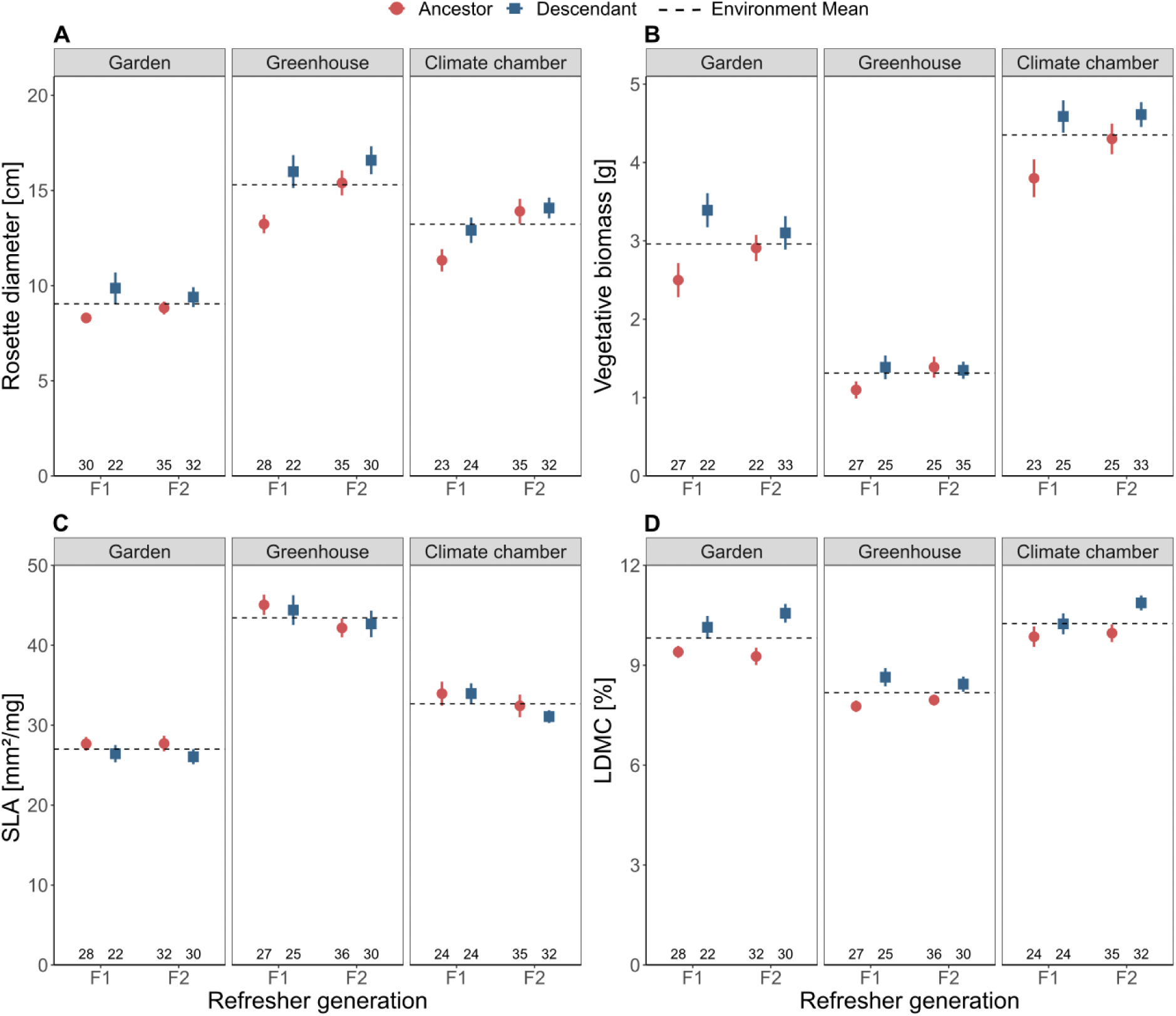
Rosette diameter (A), vegetative biomass (B), specific leaf area (C) and leaf dry matter content (D) of ancestors (blue) and descendants (red) after one intermediate generation (F1) and two intermediate generations (F2) grown in different growth facilities (garden, greenhouse, climate chamber). Shown are means and standard errors. The dotted line represents the overall mean value in each environment. Sample sizes are given at the bottom of the graph below their respective data point.

**Figure 3.**
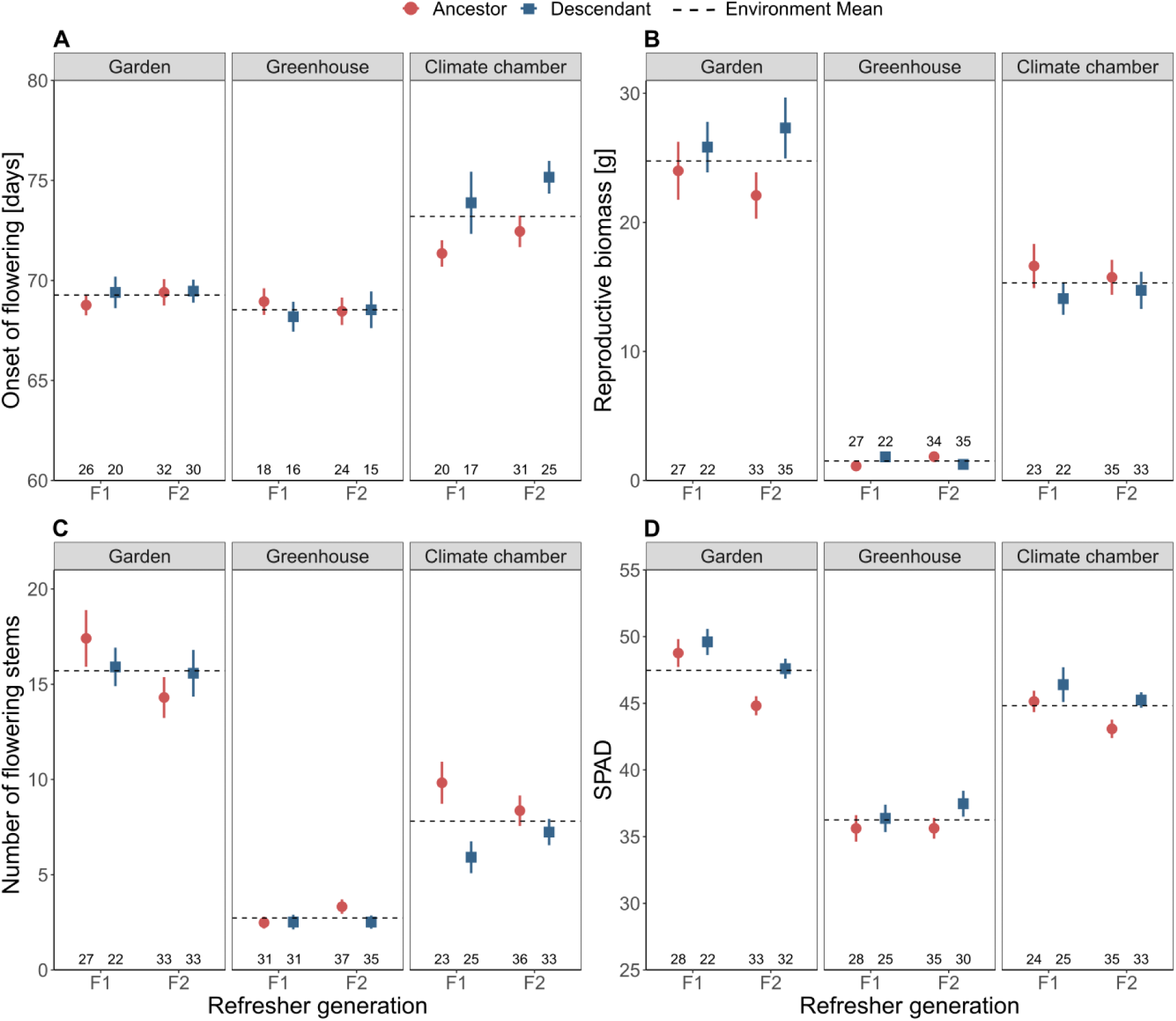
Onset of flowering (A), reproductive biomass (B), number of flowering stems (C) and SPAD values (D) of ancestors (blue) and descendants (red) after one intermediate generation (F1) and two intermediate generations (F2) grown in different growth facilities (garden, climate chamber, greenhouse). Shown are means and standard errors. The dotted line represents the overall mean value in each environment. Sample sizes are given at the bottom of the graph below their respective data point.

The temporal origin significantly affected vegetative biomass, LDMC, and onset of flowering and we found significant interactions with the growth facility (Origin × Facility) in rosette diameter and onset of flowering (Table 2). Descendants had generally 13 % higher vegetative biomass (Appendix S3A) and 8 % higher LDMC (Appendix S3B) compared to ancestors, irrespective of the growth facility or intermediate generation. Regarding the rosette diameter however, descendants had 11 % higher rosette diameter (2.1 cm) than ancestors in the greenhouse, while values were similar in the garden and in the climate chamber (Fig. 4A). Descendants also flowered 2.6 days later in the climate chamber than ancestors but flowered at similar times in the garden and greenhouse (Fig. 4B).

**Figure 4.**
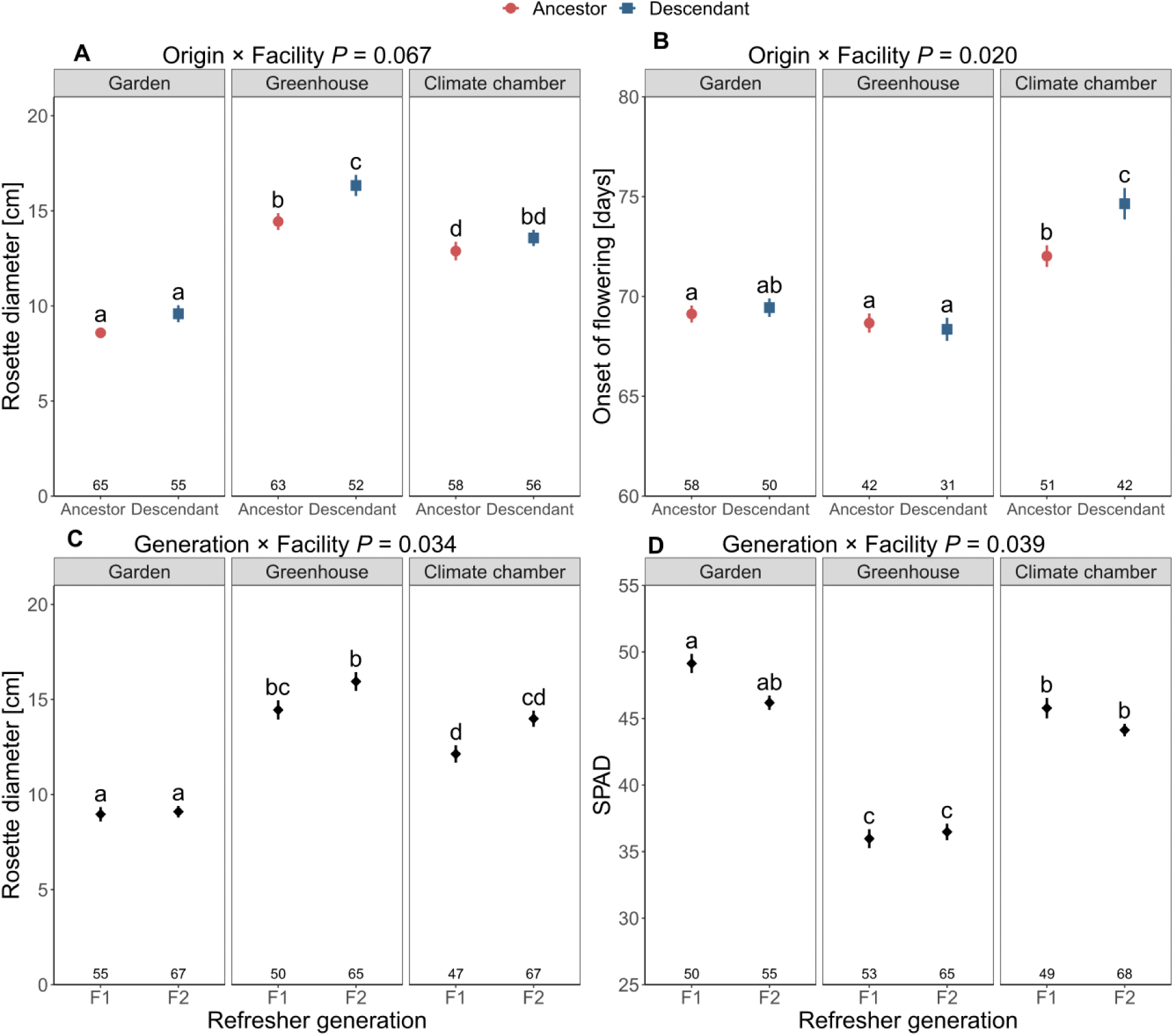
Rosette diameter (A) and onset of flowering (B) of ancestors (red) and descendants (blue) in the different environments (significant Origin × Facility effect). Rosette diameter (C) and SPAD values (D) of F1 and F2 intermediate generations in different facilities (significant Generation × Facility effect). Shown are means and standard errors. Sample sizes are given at the bottom of the graph below their respective data point.

The number of intermediate generations did not significantly affect any measured trait consistently across growth facilities, but we found significant interactions with the growth facility (Gen × Facility) in rosette diameter and SPAD values (Table 1). Although we did not find significant differences between the F1 and F2 generations within growth facilities for rosette diameter and SPAD values using Tukey post-hoc tests, the F2 had 11 % larger rosettes than the F1 both in the greenhouse and the climate chamber, while the F1 and F2 in the garden were similar (Fig. 4C). Concerning SPAD, the F1 generation showed 3 – 6 % larger values than F2 in the garden and climate chamber, while F1 and F2 did not differ in the greenhouse (Fig. 4D). We found no significant interactions between temporal origin and intermediate generation (Origin × Gen) and no significant three-way-interactions (Origin × Gen × Facility) in the measured traits (Table 2).

## Discussion

We studied the effects of three growth facilities and one *vs* two intermediate generations on the phenotypic expression of ancestor and descendant genotypes of a single population of *Leontodon hispidus*. We found very strong phenotypic differences among the three growth facilities. Furthermore, we found significant temporal origin × facility interactions in two traits, indicating that the choice of the growth facility can affect detectability of phenotypic differences. Finally, we did not find differences between intermediate generations within the growth facilities, suggesting that there is no need for multiple intermediate generations to sufficiently reduce parental and storage effects for this species.

### Differences in growth facilities

Plant responses in the different growth facilities varied greatly from each other and the observed patterns were trait specific. The outdoor garden proved to be the experimental environment where plants were most successful in their reproductive ability, as they had the most flowering stems and the highest reproductive biomass. Plants in the garden also had low SLA and high LDMC, which was probably caused by the high light availability present at that growth facilities (Table 1). It has been well established that light irradiance correlates negatively with SLA and positively with LDMC (Anten, 2005; Poorter et al., 2019). Plants in the garden also had the highest chlorophyll content (SPAD values) which could be the reason for the greater reproductive success. Previous studies also showed that leaf traits can strongly respond to water availability, with increasing dryness leading to decreasing SLA and increasing LDMC (Volaire, 2008; Poorter et al., 2009; Jung et al., 2014; Wellstein et al., 2017; Vitra et al., 2019). Accordingly, pots in the garden had the lowest soil moisture content (Fig. 1B) and high variability in soil moisture, which was caused by the exposure to natural rain events, high temperature fluctuations (Fig. 1A), and potentially higher evaporation due to wind exposure.

In the greenhouse, plants grew a large rosette diameter, had high SLA, and low LDMC. This response is in line with a strategy to increase surface area to improve light capture in low-light environments (Poorter et al., 2019). Indeed, the greenhouse, where we expected more favourable conditions than the outside garden, had the lowest light availability (Table 2) due to shading by greenhouse structural elements, but also very high temperatures (Fig. 1A). These high temperatures are also likely to contribute to the high SLA, as they facilitate cell expansion and thus, reduce leaf density and number of cell layers (Atkin et al., 1996; Poorter et al., 2009). the number of flowering stems, reproductive biomass and SPAD values were very low, indicating that even with the adjustments in leaf anatomy, the plants in the greenhouse had the lowest performance and were least successful.

Plants growing in the climate chamber had intermediate rosette diameter, SLA, SPAD values, and reproductive traits, which correlates well with the intermediate light availability (Table 1). Interestingly, plants produced the most vegetative biomass in the climate chamber out of all growth facilities. The very controlled and stable conditions might support fast vegetative growth in the climate chamber without compromising reproductive investment.

The comparisons showed significant differences among growth facilities. The main drivers of these patterns are most likely the differences in light availability and temperature, but also the variability of environmental factors could have a significant impact (Hamann et al., 2021). We expected that the garden would be the least favourable environment due to higher temperatures and stronger environmental fluctuations, leading to decreased growth and fitness of plants. However, our results indicate the opposite, which is probably due to the much higher light availability in outdoor gardens in summer compared to the greenhouse and climate chamber. Also, the garden conditions are much closer to natural conditions to which plants from the field are expected to be adapted (Lascoux et al., 2016).

### Reproducibility among growth facilities

Although it can be expected that different growth facilities cause plants to differ in their overall performance, it may also be assumed that origin and treatment effects would show qualitatively similar results across environments. Under this assumption, if a common-environment experiment would be performed in a single environment – as in most studies – the expected patterns would also be observed irrespective of this environment. Two alternative scenarios are, however, possible. First, origin or treatment effects may be observed only under specific environmental conditions and not in others. This would imply that an experiment may not always reveal the expected patterns. Second, specific origin or treatment effects may be observed under the chosen experimental conditions, but contrasting origin or treatment effects could have been observed when alternative experimental conditions would have been chosen. A consequence would be that contrasting conclusions could have been drawn, depending on the experimental environment.

In our study, descendants consistently had higher vegetative biomass and higher LDMC compared to ancestors. These results are in line with the previous findings that this population of *L. hispidus* evolved faster growth in recent decades (Karitter et al., in press), which were observed in the greenhouse in autumn. Furthermore, high LDMC has been shown to increase drought survival chances (Bongers et al., 2017; De La Riva et al., 2017). LDMC correlates well with strong cell walls and may be beneficial to maintain turgor under drought conditions (Monson and Smith, 1982). Therefore, high LDMC could have evolved in this population through selection by increasing drought events caused by climate change (IPCC, 2018). The phenotypic differences between ancestors and descendants of these two traits were consistent throughout the experimental environments, since we were able to reproduce them in the outdoor garden, greenhouse and in the climate chamber. In contrast, we found interactions of temporal origin with the experimental environment for rosette diameter and onset of flowering. Descendants had a larger rosette diameter in the greenhouse compared to ancestors, but temporal origins did not differ in any other environment. Given that the most prominent distinction of greenhouse was its low light irradiance, increasing rosette diameter may be a good strategy to increase the surface area of the leaves to capture more light. This interaction may be explained by evolution under more shaded conditions, which could have been caused by increased competition during the recent decades exacerbated by climate change (Parmesan and Hanley, 2015). At the collection site, we observed that *L. hispidus* naturally competes with grasses such as *Brachypodium pinnatum* and *Bromus erectus*, which can substantially shade this rosette species. Combined with high nutrient depositions from the atmosphere (Newman, 1995; Galloway et al., 2008; Bobbink et al., 2010) and surrounding agriculture, *L. hispidus* might have faced strong selection pressure through high competition and may have adapted its ability to plastically respond to increasingly shaded conditions. Hence, the response to high shading of the descendants was not triggered in the other environments, but this explanation needs further testing by additional experiments that include shading treatments.

We also observed later onset of flowering of descendants compared to ancestors exclusively in the climate chamber. Plants flowered generally later in the climate chamber compared to the other environments, but descendants delayed their onset of flowering substantially more than ancestors. In another resurrection study on the same population conducted in a greenhouse, descendants also flowered later than ancestors (Rauschkolb, et al., 2022), which was explained by the introduction of sheep grazing in 2007, forcing plants to flower later to escape the grazing pressure. We could not reproduce this result in the greenhouse used in this experiment and also not in the outside garden. Reason for that may be moderately different conditions in the greenhouse and also different timing of the experiment. Therefore, our results also indicate that the experimental environments used in this study provide different environmental cues for the onset of flowering. The onset of flowering is strongly dependent on environmental cues and plants can accelerate or delay it depending on local conditions in order to guarantee seed production for the next generation (Coupland, 1995). Generally, plants tend to flower later at colder temperatures (Capovilla et al., 2015), and, indeed, the average temperature was significantly lower in the climate chamber compared to the garden or greenhouse. Furthermore, the photoperiod differed in the climate chamber with approximately 2.5 h less light per day compared to the garden. Finally, the climate chamber provides the most stable conditions compared to the other environments. Descendants showed a different phenology under the shorter photoperiod, more benign temperatures and more stable conditions in the climate chamber. High variability in environmental variables in general, and even more so under climate change, is the norm under natural conditions, and perennials such as *L. hispidus* might take advantage of a stable period to grow vegetatively in order to ensure survival and increase future reproductive output, thus delaying the onset of flowering time (Tun et al., 2021). However, following this explanation, we would also expect higher vegetative biomass of descendants in the climate chamber, which was not the case. If grazing at annually recurring times selected for later flowering plants, as suggested by Rauschkolb and colleagues (2022), then the underlying process causing delayed flowering time might be through shifts in photoperiod requirement.

The majority of resurrection studies investigate the evolutionary responses of plant populations only in a single growth facility. As our study shows, depending on the trait, phenotypic differences are not guaranteed to be detected in a given experimental environment (i.e., growth facility), even if genetic differences are present. Especially when investigating evolution of plant populations, having the experimental conditions as close to their natural habitat as possible is desirable to detect evolution to the contemporary environmental conditions. With this approach, we can study the phenotypes that would occur in the field under natural conditions. However, creating deviating or even stressful environmental conditions may reveal genetic differences that are only expressed during a stressful period. The selection that caused genetic differences between ancestors and descendants could have been applied on plasticity under stressful conditions, which can only be observed under those stressful conditions. But care should be taken, because extreme conditions may also elicit responses that are not being expressed under natural extreme conditions (Ghalambor et al., 2007). When choosing between the three growth facilities we tested in this study, the best choice depends on the research aims. The garden seems to be the best option to mimic natural conditions, because of natural light, low spatial heterogeneity and contemporary weather conditions. The greenhouse seems to be the poorest choice for natural conditions, because of low light intensity and high temperatures, although it is the most used in resurrection studies (e.g., Anstett et al., 2021; Franks et al., 2007; Gay et al., 2022; Hamann et al., 2018; Kuester et al., 2016; Lambrecht et al., 2020; Nevo et al., 2012; O’Hara et al., 2021; Sultan et al., 2013; Vtipil & Sheth, 2020). Chiang and colleagues (2021) showed that environmental fluctuations can affect the phenotypic expression of multiple traits and are important to study natural-like plant growth. Here, climate chambers present an intriguing option going forward if they are programmed very closely to field conditions. Using climate or weather data from the collection sites, one could program the average temperatures, moisture, light spectrum and their daily to seasonal variability to have the conditions very close to the field, while still having a high level of control (Poorter et al., 2016). This idea has already been successfully applied by Heuermann and colleagues (2023) who managed to simulate whole seasons in a reproducible manner in their specialized indoor growth facility. Additionally, extreme events can be modelled (e.g., heatwaves) as treatments to further investigate if differences are also expressed under such conditions, especially if these conditions are potential selection agents. Very controlled and homogenous conditions on the other hand, can be useful if only genetic differences per se are of interest and not how they relate to the actual natural conditions. Especially low temporal variation in experimental conditions is important to provide similar conditions throughout all ontogenetic stages, and to avoid interactions of ontology and the environmental conditions. Furthermore, it is possible to infer adaptive traits in the field from genotypes grown indoors by using modelling of the environmental conditions (Bouidghaghen et al., 2023).

### Intermediate generations

We found no differences between the two intermediate generations used in this study within each of the growth facilities, indicating either that no significant parental effects were present beforehand at all or that they were removed after the first intermediate generation. The magnitude and nature of parental effects can be strongly dependent on the environmental stresses experienced by the parents, as Latzel and colleagues (2023) showed that theses stresses can affect the fitness of the offspring by up to 35 % in *Arabidopsis thaliana*. It is likely that environmental stresses differed between ancestors and descendants of our study population, as climate change increased the frequency and duration of droughts and heatwaves (IPCC, 2018). However, if there were any detectable parental effects, these have been eliminated in the first intermediate generation, which has also been found in other studies (Agrawal, 2002; Gianoli, 2002). Since we did not have seed material of the originally collected sample – i.e. before the intermediate generations –, we cannot quantify how much the first intermediate generation reduced parental and storage effects. Thus, we cannot know whether there were detectable parental effects in the first place. Theoretically, long-term storage of seeds may cause carry-over effects into the F1-generation in suboptimal storage conditions, with resurrected plants having lower fitness due to the storage and producing lower quality seeds (Gebeyehu, 2020). The seed material used in this study was stored at –20°C after drying at 15% RH which are optimal conditions to ensure viability for several decades (Solberg et al., 2020). The implications of potential carry-over effects can of course be very dependent on the storage condition and storage length, and other species might be affected more than *L. hispidus*. Thus, multispecies experiments are needed to advance our understanding of parental and storage effects and to make informed choices regarding the amount of required intermediate generations for a given species.

## Conclusions

Our study shows that the choice of the growth facility in common-environment experiments can potentially impact the expression of phenotypic differences among genotypes, thereby affecting the conclusions. Thus, studying evolution of plant populations in only a single environment might result in incomplete or even deficient interpretations for some traits. Hence, it is important to carefully choose the growth facility or even use multiple facilities. Outdoor garden experiments might be a good and simple option with regard to studying rapid evolution as plants will experience more natural conditions and the contemporary climate which they are expected to have evolved to. However, if environmental variables from the population are well known, using climate chambers might be a good alternative with a high level of control and detailed programming to encompass realistic natural conditions as well as less or more extreme conditions that may occur under natural conditions, e.g., as additional treatments. Finally, growing a second intermediate generation rather than only one intermediate generation did not impact the genetic differences of ancestors and descendants within growth facilities, suggesting that only one intermediate generation would be sufficient to reduce detectable parental and storage effects, if there were any in the first place. Overall, future studies should be aware of implications regarding reproducibility and wisely choose the experimental conditions.

## Acknowledgements

The authors thank the Deutsche Bundesstiftung Umwelt (DBU) for their financial support by a PhD scholarship (20020/678) to PK. We thank Lutz Stübing, Susanne Pietsch, Robert Anton and the gardeners from the “Wissenschaftsgarten” of Goethe University Frankfurt for their support, and Charlotte Møller for her help with the data loggers.

## Data archiving

The data that support the findings of this study will be made available at Dryad after this manuscript is accepted for publication.

## Supporting information

**Appendix S1.**
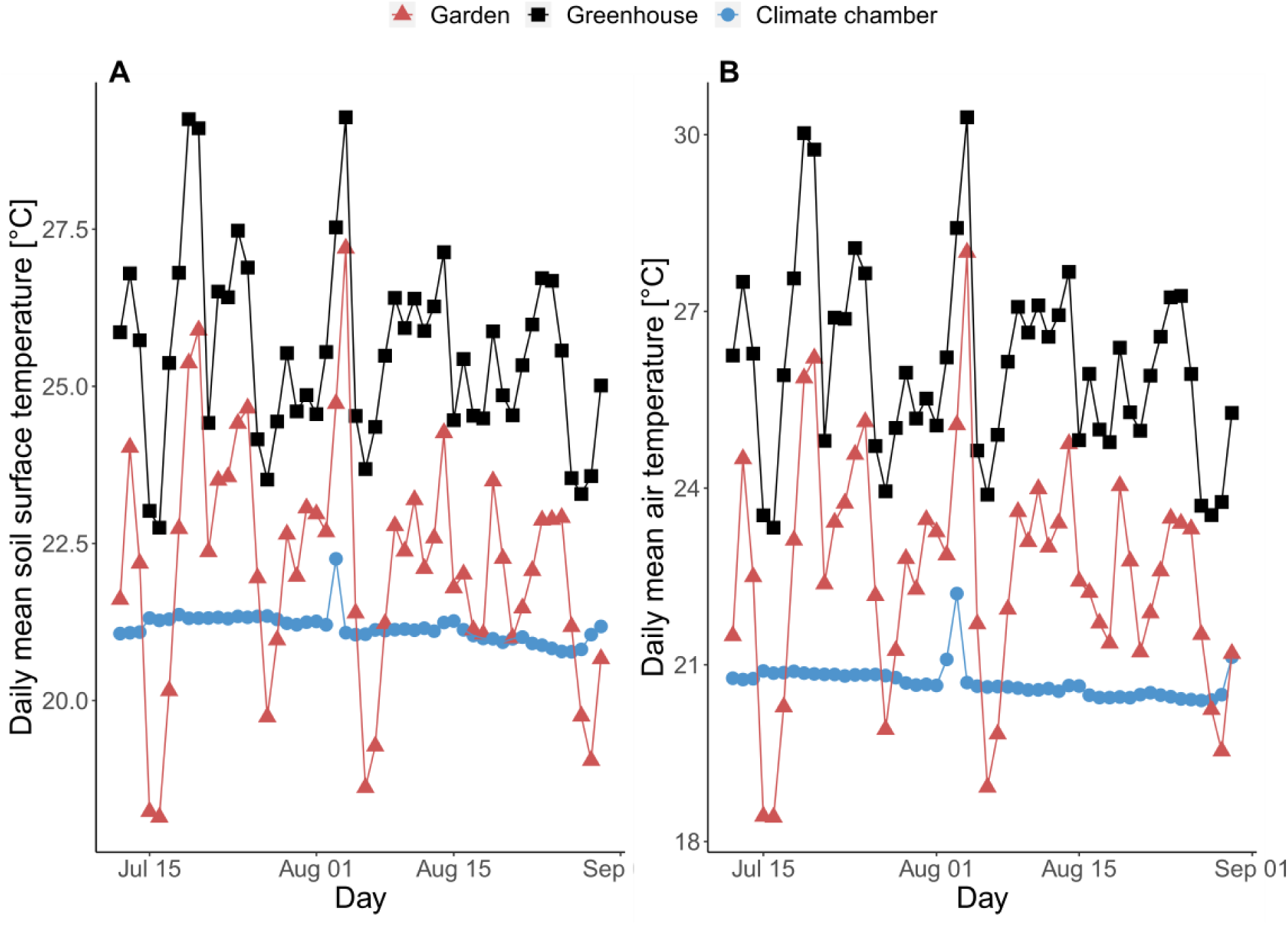
Daily mean soil surface temperature (A) and daily mean air temperature (B) of four random pots grown in different growth facilities. The growth facilities are garden (red triangles), greenhouse (black squares), and climate chamber (blue circles).

**Appendix S2.** Results before adjustments of the statistical models testing the effects of temporal origin (ancestors, descendants), intermediate generation (F1, 2), growth facility (garden, greenhouse, climate chamber) and their interactions on the response variables (y) rosette diameter, vegetative biomass, specific leaf rea (SLA), leaf dry matter content (LDMC), onset of flowering, reproductive biomass, number of stems and SPAD measurements. We used linear mixed effects odels with initial size as covariate and maternal line as random factor followed by ANOVA’s. Significant values (*P* < 0.05) are shown in bold.

|  |  | **Rosette**<br>**diameter** | **Vegetative**<br>**biomass** | **SLA** | **LDMC** | **Onset of**<br>**flowering** | **Reproductive**<br>**biomass** | **Number of**<br>**stems** | **SPAD** |
| --- | --- | --- | --- | --- | --- | --- | --- | --- | --- |
|  | *df* | *P* value | *P* value | *P* value | *P* value | *P* value | *P* value | *P* value | *P* value |
| **Initial size** |  | **<0.001** | **<0.001** | **0.004** | **<0.001** | **<0.001** | 0.15 | 0.708 | 0.379 |
| **Origin** | 1 | 0.118 | **0.007** | 0.676 | **0.005** | **0.014** | 0.969 | 0.210 | 0.101 |
| **Gen** | 1 | 0.145 | 0.414 | 0.204 | 0.639 | 0.198 | 0.673 | 0.623 | 0.130 |
| **Env** | 2 | **<0.001** | **<0.001** | **<0.001** | **<0.001** | **<0.001** | **<0.001** | **<0.001** | **<0.001** |
| **Origin ×**<br>**Gen** | 1 | 0.584 | 0.106 | 0.633 | 0.264 | 0.408 | 0.659 | 0.299 | 0.339 |
| **Origin ×**<br>**Env** | 2 | **0.021** | 0.214 | 0.593 | 0.590 | **0.004** | 0.128 | 0.068 | 0.899 |
| **Gen × Env** | 2 | **0.009** | 0.678 | 0.371 | 0.507 | 0.661 | 0.912 | 0.140 | **0.011** |
| **Origin ×**<br>**Gen × Env** | 2 | 0.902 | 0.728 | 0.785 | 0.219 | 0.740 | 0.232 | 0.176 | 0.817 |

**Appendix S3.**
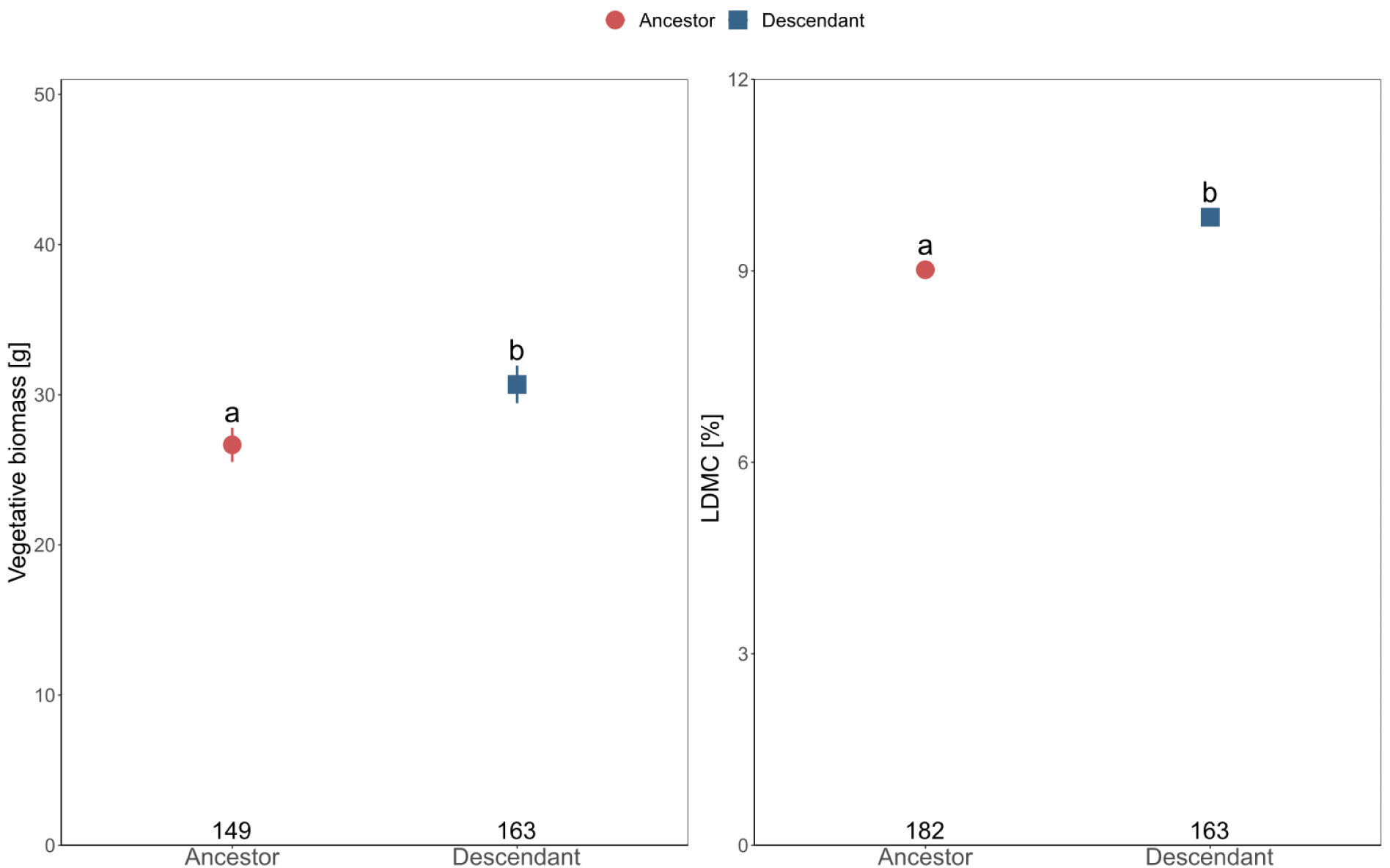
Vegetative biomass (A) and LDMC (B) of ancestors and descendants (significant Origin effect). Shown are means and standard errors. Standard errors of LDMC are too small to be visible. Sample sizes are given at the bottom of the graph below their respective data point.

## Notes

### Competing Interest Statement

The authors have declared no competing interest.

